# Density-Weighted Concentric Ring Trajectory using simultaneous multi-band acceleration: 3D Metabolite-cycled Magnetic Resonance Spectroscopy Imaging at 3 T

**DOI:** 10.1101/628594

**Authors:** Pingyu Xia, Mark Chiew, Xiaopeng Zhou, Albert Thomas, Ulrike Dydak, Uzay E Emir

## Abstract

Simultaneous multi-slice or multi-band parallel imaging is frequently used in fMRI to accelerate acquisition. In this study, we propose a new a non-water-suppressed multi-band MRSI using a density weighted concentric ring trajectory (DW-CRT) acquisition. The properties of multi-band acquisition scheme were compared against single-band acquisition in phantoms and in-vivo experiments. High-quality spectra were acquired for all simultaneously acquired three slices from a limited volume of interest with an in-plane resolution of 5 × 5 mm2 in 19.2 min. The achieved in vivo spectral quality of the multi-band method allowed the reliable metabolic mapping of four major metabolites with Cramér-Rao lower bounds below 50%, using a LCModel analysis. Metabolite maps and LCModel quality metrics were compared between multi-band and single-band acquisition schemes for both phantom and in-vivo measurements. Structural similarity (SSIM) index and conventional coefficient of variance analysis of both phantom and in-vivo measurements revealed a SSIM index higher than 0.83, corresponding to a coefficient of variance of less than 30%. These findings demonstrate that our multi-band DW-CRT MRSI demonstrate a multi-band acceleration, simultaneously acquired 3 slices, with no apparently loss in spectral quality, reducing scan time by a factor of 3 without any significant penalty.

## Introduction

Magnetic resonance spectroscopic imaging (MRSI) measures neurochemical profiles over larger regions non-invasively. However, the practicality of MRSI methods is compromised by low concentrations of metabolites relative to water, leading to low signal-to-noise ratios (SNRs) as well as long acquisition times due to spectral and spatial encoding(1).

There have been several methods proposed to accelerate acquisition and reduce imaging durations for MRSI. Accelerated k-space trajectories such as EPSI(2,3), spiral (4), concentric rings(5) and rosette(6) showed significant speed-ups for MRSI. Besides, several parallel imaging methods have been applied to MRSI to for use with conventional (7) and accelerated k-space trajectories (8). Recently, a feasibility study was performed to evaluate the use of CAIPIRINHA to accelerate MRSI along three spatial dimensions (9). Simultaneous multi-slice (SMS) or multi-band (MB) parallel imaging is frequently used in fMRI to accelerate acquisition (10), and recently a feasibility study using MB has been performed to accelerate MRSI acquisition along slice selection direction using EPSI (11).

The intent of this work, therefore, is to develop a non-water-suppressed MB MRSI using a density weighted concentric ring trajectory (DW-CRT) acquisition. Sampling k-space using a DW pattern to shape the spatial response function improve side lobe artefacts and signal-to-noise ratio (SNR) (12). A non-water-suppressed metabolite-cycling is proposed for simultaneous detection of the metabolites and water signals at short echo time (TE = 14 ms) using stimulated echo acquisition mode (STEAM) localization (13). This provides the required information for voxel-wise single-scan frequency, phase, gradient-induced sideband and eddy current correction. In this study, we demonstrate a MB acceleration, simultaneously acquired 3 slices, with no apparently loss in spectral quality, reducing scan time by a factor of 3 without any significant penalty. To validate metabolite profiles obtained using the newly developed acquisition technique, we compare profiles quantified from single-band acquisition from the same slices.

## Methods

### Validation study

This study is approved by the Institutional Review Board at Purdue University. Five healthy volunteers (3 females, aged 22.6± 1.67 year) participated. All participants gave their written informed consent prior to participation.

### Non-water suppressed metabolite cycling MRSI acquisition

Non-water-suppressed metabolite-cycling MRSI was acquired using parameters described in Emir et al (13,14). Briefly, the metabolite-cycling MRSI acquisition was achieved by the inversion of the upfield and downfield (N_directions_=2) spectral resonances before localization pulses with a gap of 9.6 ms so that water spectra (addition of both spectra) and metabolite spectra (subtraction of both spectra) can be kept simultaneously. The water spectra would later be used for frequency and phase alignment, eddy current correction, and as the internal concentration reference. The inversion pulse is the combination of hyperbolic secant pulse HS_l/2_ and tanh/tan pulse [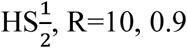, Tp; tanh/tan, R=100, 0.1 Tp], providing a sharp transition band of 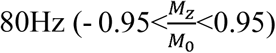 and a broad inversion band of 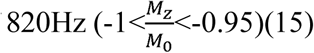.

### Multi-band STEAM Localization

In slice-accelerated MB imaging, the number of simultaneously excited slices dictates the number of bands required in the radio frequency (RF) pulse. Because multiple slice-selective pulses are applied simultaneously, the multi-band approach quickly runs into a peak power limitation. Thus, we used the optimal phase modulation method to reduce multi-band pulse peak power as proposed by Wong et al, resulting in a 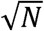 (instead of N) times of increase of peak *B*_l_ compared with that of single band pulse (16). The peak *B*_l_ for 4 ms single band pulse is 8.22 μT, while that of 4-ms multi-band pulse is 18.38 μT. The spectral bandwidth for each band is 1.55KHz for both single band and multi-band pulses. Figure 1a illustrates the phase-modulated multi-band RF pulse used for multi-band excitation.

**Figure 1.**
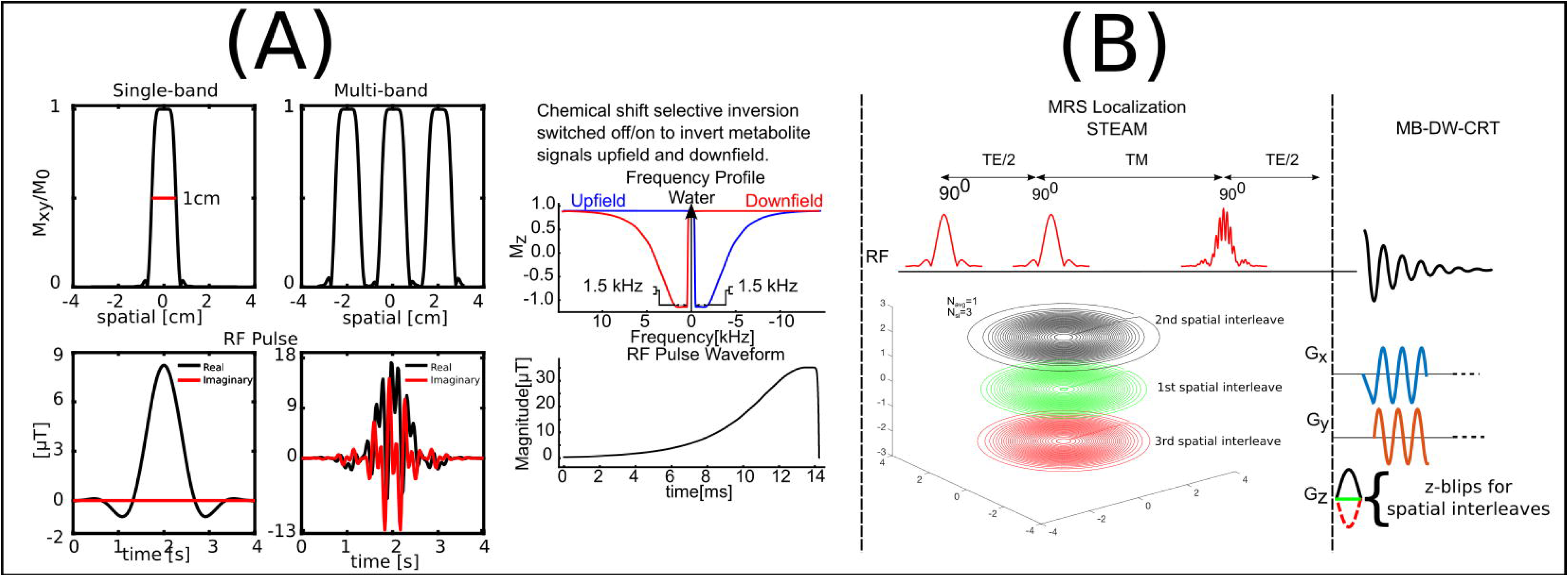
(A) RF pulse and corresponding slice excitation profiles of single band and multiband pulses. The phase modulation of multi-band pulse reduced the peak ***B***_l_ while kept the bandwidth of each band and pulse duration the same. (B) The proposed MB-DW-CRT MRSI method without water suppression, using the third STEAM pulse as multi-band excitation pulse. 96 rings with different radii are distributed evenly in three set of rings with different z-blip gradients. Single-slice sequence has the same 96 rings in kx-ky plane without encoding with z-blip gradients.

### Z-blipped Density Weighted Concentric Ring Trajectory

The k_x_-k_y_ trajectory was composed of 96 distinct concentric rings with 160 equidistant points on each ring. Within one repetition time (TR) duration, only one ring will be acquired and repeated 512 times for spectral encoding. A DW method is used to determine the radii of the k-space rings because the non-uniform k-space sampling density allows effective windowing during acquisition instead of post-processing, providing higher efficiency (17). A Hanning-window sampling density is selected due to its good compromise between sensitivity and localization (18), and is given by:

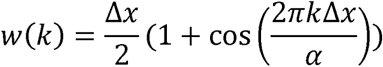

where Δ*x* is the nominal spatial resolution, and *α* is the parameter adjusting the effective spatial resolution by modulating the width of the spatial response function (SRF) main lobe. The parameter α is set to 1 for the maximum radius of 1/(2Δx).

For multi-band acquisition, three 10 mm slices were excited simultaneously with a 20 mm distance between center of adjacent slices. To decrease the coil geometry factor, a blipped-CAIPI(19) application of z-gradient blips were applied before the start of each ring of DW-CRT (Figure 1b). The corresponding k_z_ were 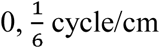, and 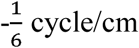, repeating every three rings. This results in three spatial interleaves (number of spatial interleaves, N_si_ = 3) of 32 DW concentric rings (number of rings, N_ring_ = 32) with different phase modulation and aliasing patterns in three slices. The middle slice remained unmodulated, the top slice experienced an additional (0, 120, −120) phase cycle, and the bottom slice experienced an additional (0, −120, 120) phase cycle of every three rings. Unlike the purely spatial shift in phase encoding direction in CAIPIRINHA(20) and blipped-CAIPI EPI, the aliasing patterns of top and bottom slices are spreading out because the Fourier transform of the phase modulation is no longer a spatial Dirac function (21).

### MRI data acquisition

Phantom and in-vivo scans were acquired using a Siemens Prisma 3-Tesla (Siemens, Germany) scanner and a 20-channel head array receive coil. T1-weighted MP-RAGE images (TR = 2200 ms, TE = 2.48 ms, TI = 900 ms, flip angle = 8°, 176 transverse slices, 0.9 × 0.9 × 1mm^3^) were acquired for MRSI VOI placement. GRESHIM (gradient-echo shimming) was used for B_0_ shimming(22).

### Phantom acquisitions

At the beginning of all experiments, single-band MRSI data were acquired separately from the same locations as the multi-band MRSI for calculation of coil sensitivities and quality comparison analysis between multi-band and single-band analysis. The single-band acquisition used the same acquisition scheme as multi-band without the z-gradient blips and the multi-band excitation pulse. The multi-band MRSI acquisition was tested on two phantoms. The first phantom was the American College of Radiology (ACR) structural phantom. The second phantom measurement was from a multi-compartment MRS phantom containing six hollow spheres filled with different concentrations of NAA (with a maximum concentration of 15 mmol) and Creatine (Cr, with a maximum concentration of 15 mmol). The NAA and Cr distributions measured by single-band and multi-band acquisition schemes were used to evaluate multi-band sequence performance. The FOV of the MRSI acquisition was 240 mm × 240 mm. The STEAM localization (TR =1 s, TE = 14 ms, and a mixing time (TM) = 32 ms) was 160 mm × 160 × 10 mm. To cover the 48 × 48 grid, 32 DW concentric rings resulting in an individual voxel size of 0.25 mL were acquired with three spatial interleaves for single band and three z-gradient blips for multi-band acquisition. The multi-band acquisition was acquired with six averages (Navg=6) resulted in a total acquisition duration of 19.2 min (Navg × Nrings × Nsi × T_R_ × Ndirections=1152 s) whereas single band acquisition of three slices with one average was completed in 9.6 min (Nslc × Nrings × Nsi × T_R_ × Ndirections = 576 s).

### In vivo Acquisition

In-vivo acquisition was performed with identical parameters, where only the in-plane dimension of the STEAM localization was reduced to 80 mm × 110 mm so that only the top slice extended to the subcutaneous lipid layer while the middle slice and bottom slice covered as large a brain region as possible.

### Post-processing

Reconstruction algorithms were implemented in MATLAB (MathWorks, Natick, MA, USA). Gridding reconstruction was performed without using any post-hoc density compensation because the density weighting method relies on the different sampling density of different k-space locations. Sensitivity maps for SENSE reconstruction were estimated from the single-band calibration data based on the adaptive filtering method to optimize the SNR(23). The NUFFT(24) and SMS(21) toolboxes were used to perform the non-cartesian SENSE reconstruction of single band and multi-band MRSI data.

After SENSE reconstruction of MRSI data, upfield and downfield single-shot metabolite-cycled FIDs in each voxel were frequency and phase corrected. Then, they were summed to generate water spectra and subtracted to get metabolite spectra(13). The residual water peak of metabolite spectra was removed using the Hankel-Lanczos singular value decomposition (HLSVD) algorithm(25).

Lipid removal was performed on all three slices using a lipid-basis penalty algorithm. The lipid basis was generated from the top slice subcutaneous lipid area, the mask of which is hand-drawn based on the observed contrast between the brain and non-brain tissue in the water MRSI images. An L2-penalty iterative reconstruction with a regularization parameter of 10^−l^ was applied to remove lipid signals from metabolite MRSI assuming that metabolite spectra from brain and lipid spectra from lipid mask were orthogonal(26). The voxels processed within the brain region were then passed to metabolite analysis.

The LCModel package was used to quantify the metabolite spectrum for each MRSI voxel. The model spectra of alanine (Ala), aspartate (Asp), ascorbate/vitamin C (Asc), glycerophosphocholine (GPC), phosphocholine (PC), Cr, phosphocreatine (PCr), GABA, glucose, glutamine (Gln), glutamate (Glu), glutathione(GSH), lactate (Lac), myo-Inositol (myo-Ins), NAA, N-acetylaspartylglutamate (NAAG), and taurine(Tau) were generated based on previously reported chemical shifts and coupling constants by the GAMMA/PyGAMMA simulation library of VeSPA (Versatile Simulation, Pulses and Analysis) according to a density matrix formalism. Simulations were performed using the same RF pulses and sequence timings as those on the 3 T system in use. Five LCModel-simulated macromolecule resonances were included in the analysis at the following spectral positions: 0.91, 1.21, 1.43, 1.67, and 1.95ppm. LCModel analysis was performed in a spectral range between 0.5 and 4.2 ppm. Concentrations were calculated using the unsuppressed water spectrum as an internal reference.

### g-Factor analysis

The pseudo multiple replica method is used to simulate image noise amplification due to the computational infeasibility of direct calculation (27). In our case, it is because the z-blipped concentric ring k-space trajectory doesn’t yield discrete aliased voxels during SENSE reconstruction. The noise covariance matrix between all channels was calculated from complex noise acquired with all RF pulses turned off. Multiple pseudo replicas were simulated by adding random gaussian noise synthesized based on the estimated coil noise covariance matrix. Image noise is estimated from standard deviation map of reconstruction results of those noise replica. The g-factor map is then quantified dividing multi-band image noise by single band image noise. 1000 pseudo replica were simulated for both multi-band and single band sequences.

### Structural Similarity Index (SSIM) analysis

In this study, we explore the use of the Structural Similarity Index (SSIM) to measure the similarity of generated water and metabolite images of multi-band acquisition against the single band ones. Conventional voxel-wise coefficient of variances (CV) between multi-band and single band water images and metabolite maps were also calculated to assess the deviation of multi-band from single band data.

## Results

### Slice unaliasing evaluation of multi-band MRSI acquisition

The performance of slice unaliasing of multi-band MRSI acquisition was evaluated based on the reconstructed water images (magnitude of first-time point of FID in each voxel) of ACR phantom as illustrated in Figure 2. The left panel of Figure 2 shows the g-factor map of the proposed acquisition and reconstruction steps. The proposed acquisition scheme and reconstruction pipeline allowed a mean g-factor value of 1.096. Although the final resolution of the images generated from FIDs of multi- and single-band acquisitions was poorer than the MP - RAGE images (0.25 mL versus 0.001 mL), the non-water-suppressed metabolite-cycling MRSI and its reconstruction generated a spectroscopic image with structural information similar to that of MPRAGE. Residual artifacts originating from proposed unaliasing procedure mainly appeared near the edge, and they were not evident in inner regions. Distinct components of each slice are not visibly contaminating other slices, such as the grids, wedges with different lengths, and hole arrays, demonstrating the capability of SENSE unaliasing of blipped concentric ring trajectories.

**Figure 2.**
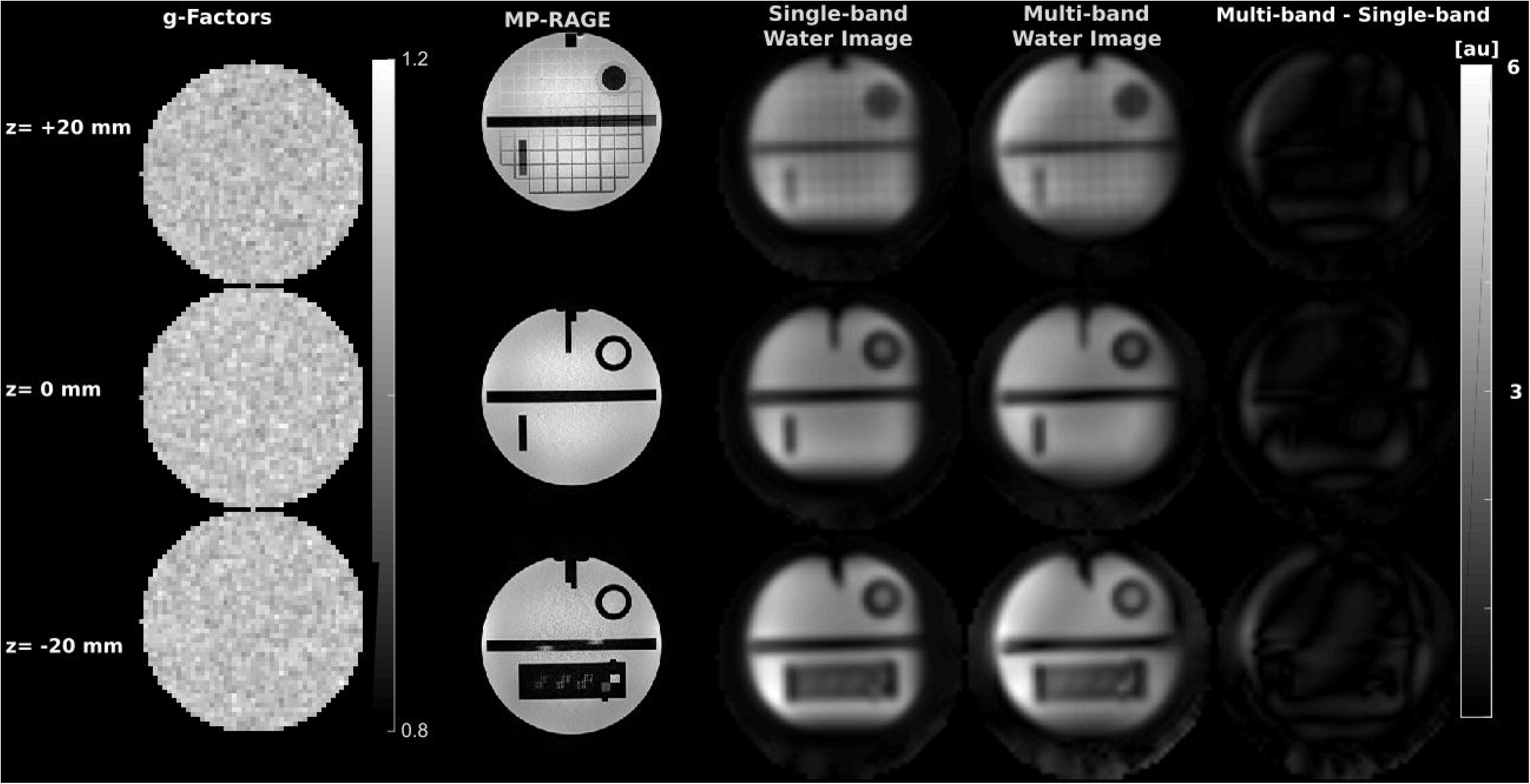
g-factor maps and Slice unaliasing performance of ACR phantom acquisitions. Three slices were positioned deliberately to include distinct and delicate features. Multi-band MRSI water images replicate corresponding single band images accurately in most regions without obvious interslice leakage. The spectroscopic water images were interpolated to a 256□×□256 matrix for display.

### Metabolites quantification accuracy of multi-band MRSI acquisition – Phantom MRSI

Figure 3 shows results from the multi-compartment MRS phantom. Figure 3 shows representative spectra from single and multi-band in a 3 × 3 grid marked on the reconstructed water images. The accuracy of metabolites quantification of multi-band MRSI acquisition was estimated based on the comparison against the single-band MRSI results. Figure 4 shows MP-RAGE, water, NAA, Cr and corresponding LCModel quality metrics, SNR, and FWHM maps. The multi-band SNR has been divided by the square root of the Navg of multi-band, 6, to compensate the Navg difference between single-band and multi-band acquisitions. The similarity between generated metabolites images is visually distinct. SSIM and CV analyses further corroborated this similarity. As shown in Figure 5, the SSIM maps of tNAA maps from three slices demonstrated very high similarity indexes in the compartments where NAA presents. The mean SSIM of NAA in the hollow spheres was higher than 0.83, corresponding to a coefficient of variance of less than 30%. However, Cr maps showed homogeneous distributions of SSIM in three slices since the presence of Cr in the container. Similarly, FWHM and SNR maps also have high SSIM values, indicating the minimal spectra quality degradation caused by multi-band acceleration. The relationship between SSIM and CV is illustrated in Figure 5. The SSIM values corresponding to a CV of 30 are marked on the plot. The numbers of voxels for each slice with a CV of less than 30 % were listed in Supplementary Table 1.

**Figure 3.**
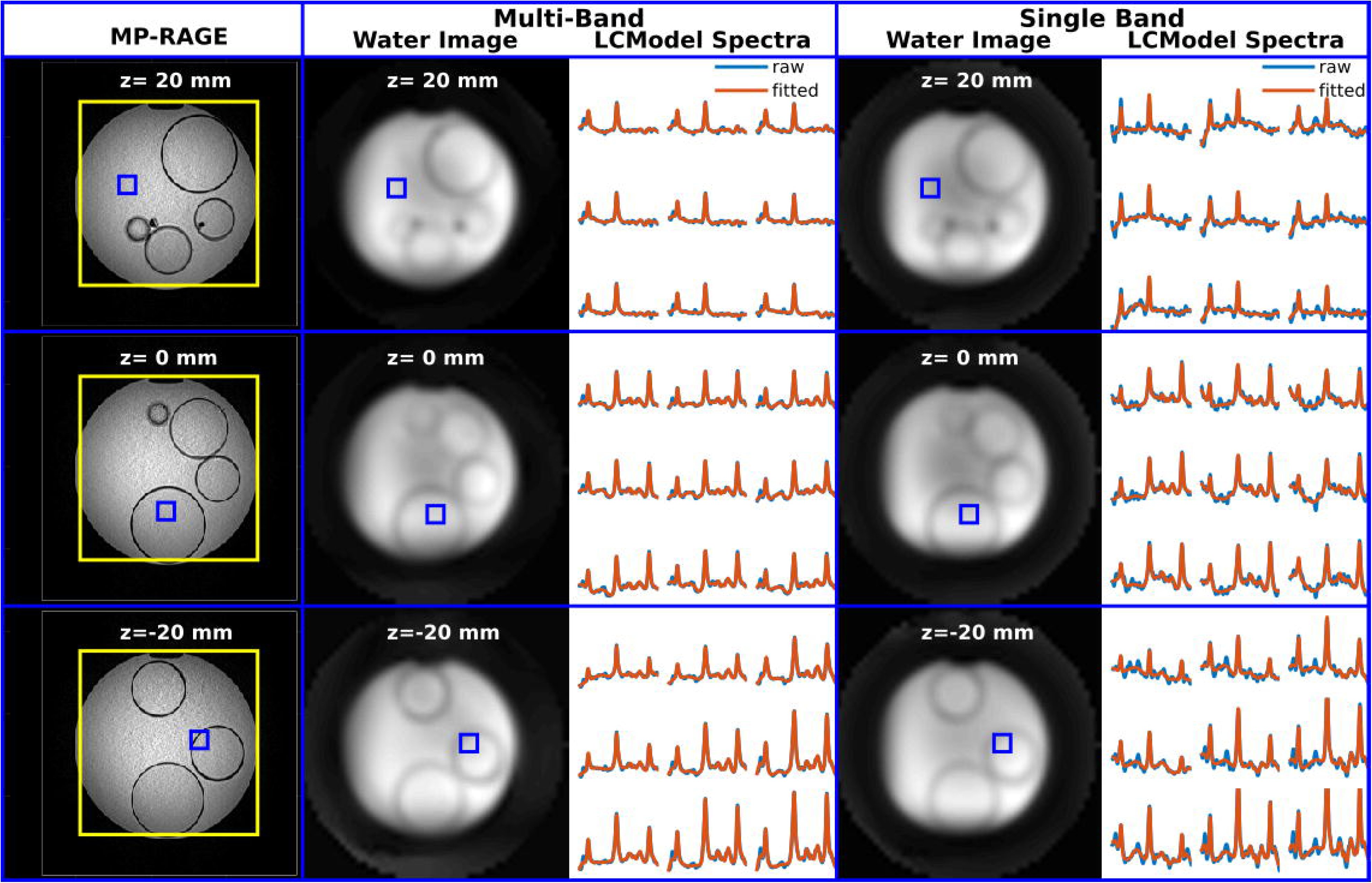
MP-RAGE images, multi-band and single band water images, and 3×3 spectra from positions marked by blue box. The blue box of the top slice (z = 20 mm) is placed outside small spheres, where only Cr exist. The blue box of the middle slice (z = 0 mm) is placed inside one of the small sphere, where metabolite distribution should be homogeneous. The blue box of the bottom slice (z = −20 mm) is placed at the edge of one small sphere, where metabolite distribution should change from inside to outside. Spectra were illustrated in a spectral range between 1.7 and 4.2 ppm. The spectroscopic water images were interpolated to a 256□×□256 matrix for display.

**Figure 4.**
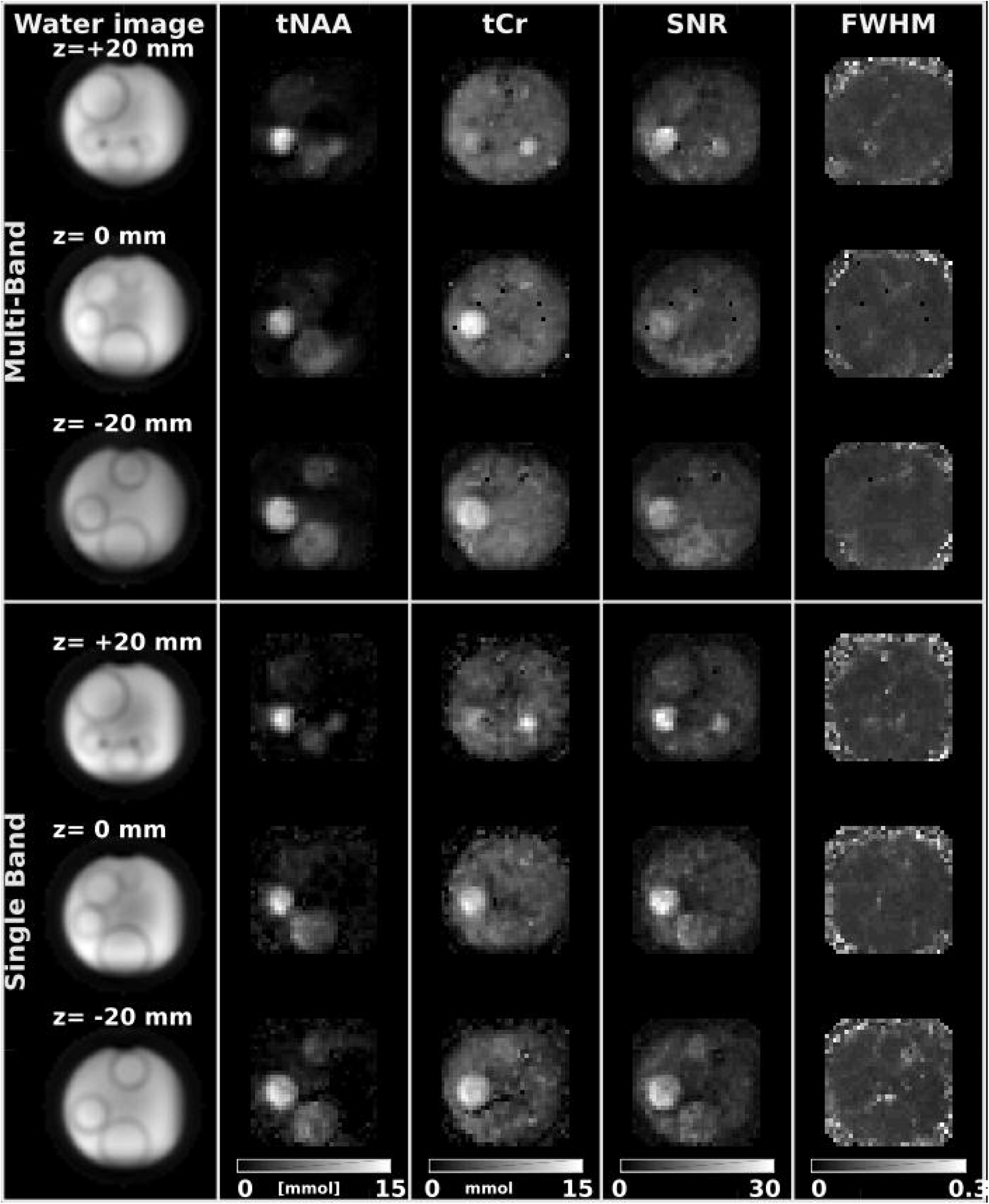
Water, tNAA, tCr, SNR, and FWHM of multi-band (upper) and single-band(lower) images. Single-band acquisition looks noisier since the Navg was 1 while Navg of multi-band acquisition is 6. The SNR of multi-band was divided by √6 to ensure the comparison between multi-band and single-band results was independent from Navg. The spectroscopic water images were interpolated to a 256□×□256 matrix for display.

**Figure 5.**
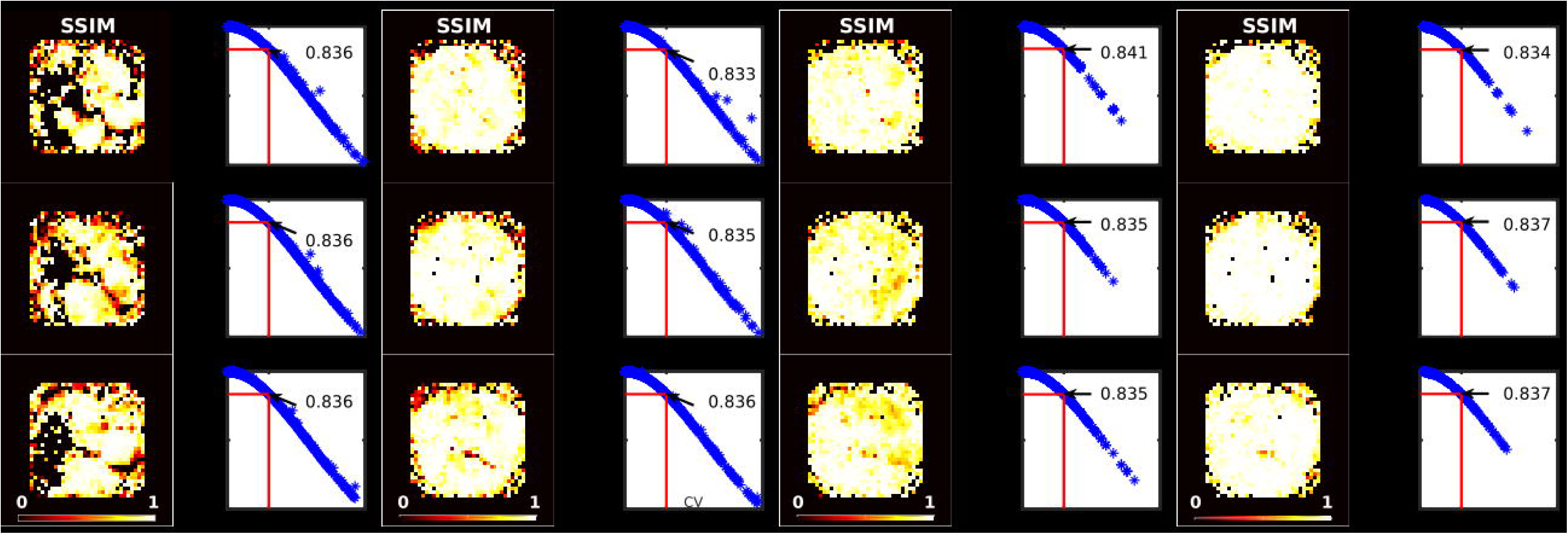
Structural similarity (SSIM) quality maps between multi-band and single-band acquisitions were were illustrated for tNAA, tCr, SNR and FWHM as estimated by LCModel. Comparison of the SSIM index and coefficient of variance (CV) between multi-and single-band acquisitions were also illustrated. A CV of 30 % and its corresponding SSIM index for each map were highlighted with a black arrow.

### Metabolites quantification accuracy of multi-band MRSI acquisition – in vivo MRSI

Figure 6 shows the representative single- and multi-band spectra from a volunteer. As illustrated on a 3 × 3 grid from three slices, metabolite spectra with LCModel fits from multi-band MRSI resulted in similar spectral features of single-band MRSI. For example, ventricular CSF showed low levels of all metabolite signals (slice z=-20mm), and white matter spectra showed an increase in tCho/tCr ratio compared to gray matter spectra (slice z=0).

**Figure 6.**
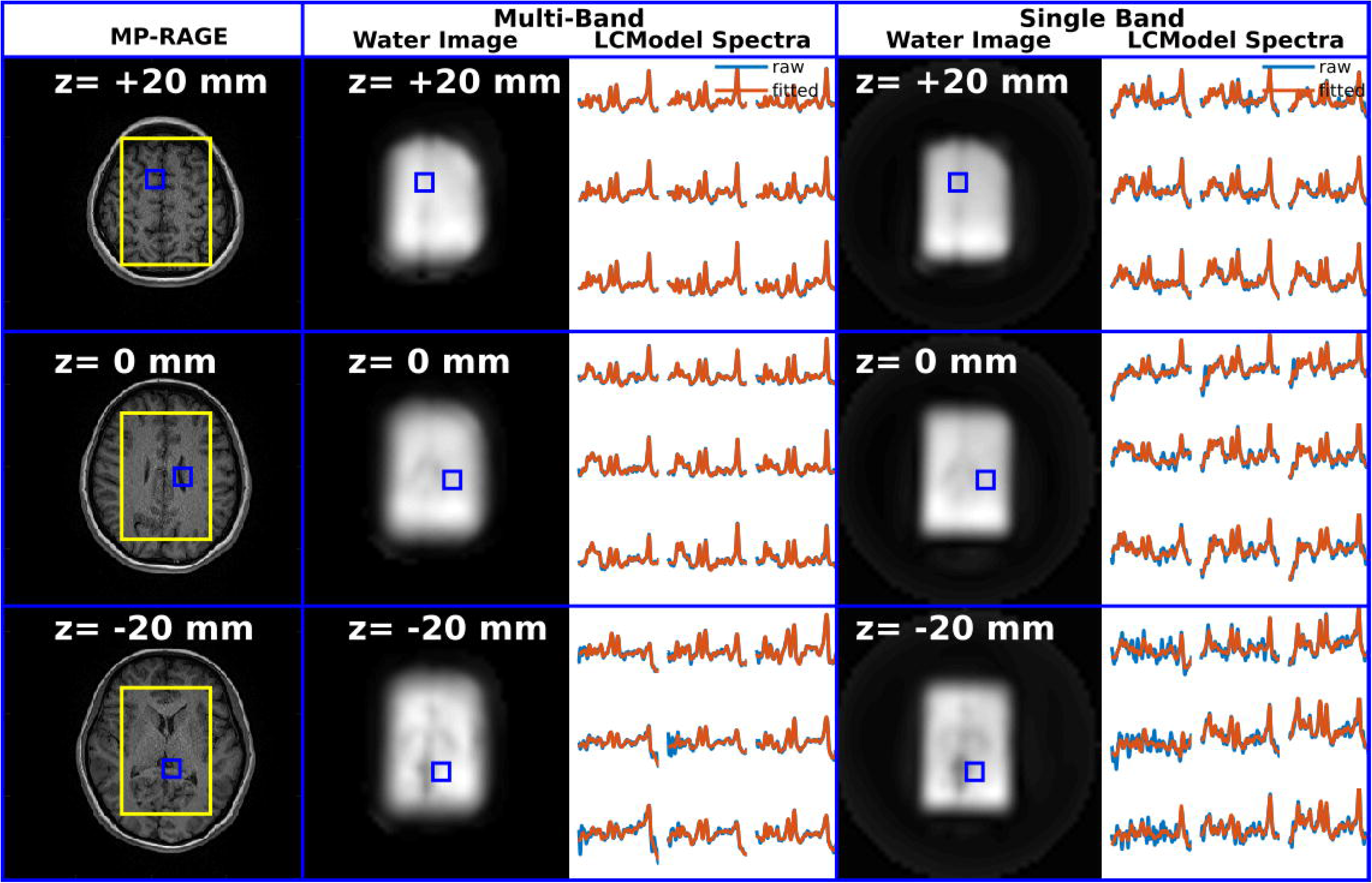
MP-RAGE, multi-band and single band water images, and 3×3 spectra from positions marked by blue box. Spectra were illustrated in a spectral range between 1.7 and 4.2 ppm. The spectroscopic water images were interpolated to a 256□×□256 matrix for display.

For the multi-band MRSI acquisition with a resolution of 0.25 mL, the achieved in vivo spectral quality allowed reliable quantification of major brain metabolites with a CRLB of less than 50% using LCModel analysis (Figure 7). Concentration distributions of the metabolites quantified were in good agreement with the single-band acquisition and previously reported values acquired from the same brain locations, and revealed significant variations between different brain tissues. The lack of metabolite signals in the region of ventricles resulted in low concentration estimation for both single- and multi-band tNAA, tCr, and Glx maps of the bottom slices (z=-20 mm). The contrast of gray matter and white matter was also evident on tCr and Glx maps of the superior slices (z = 0 and +20 mm). In agreement with the phantom experiment, SSIM and CV analyses resulted in similar findings (Figure 8). Most voxels in three slices resulted in SSIM value higher than 0.83 corresponding to a CV of 30%, Supplementary Table 2.

**Figure 7.**
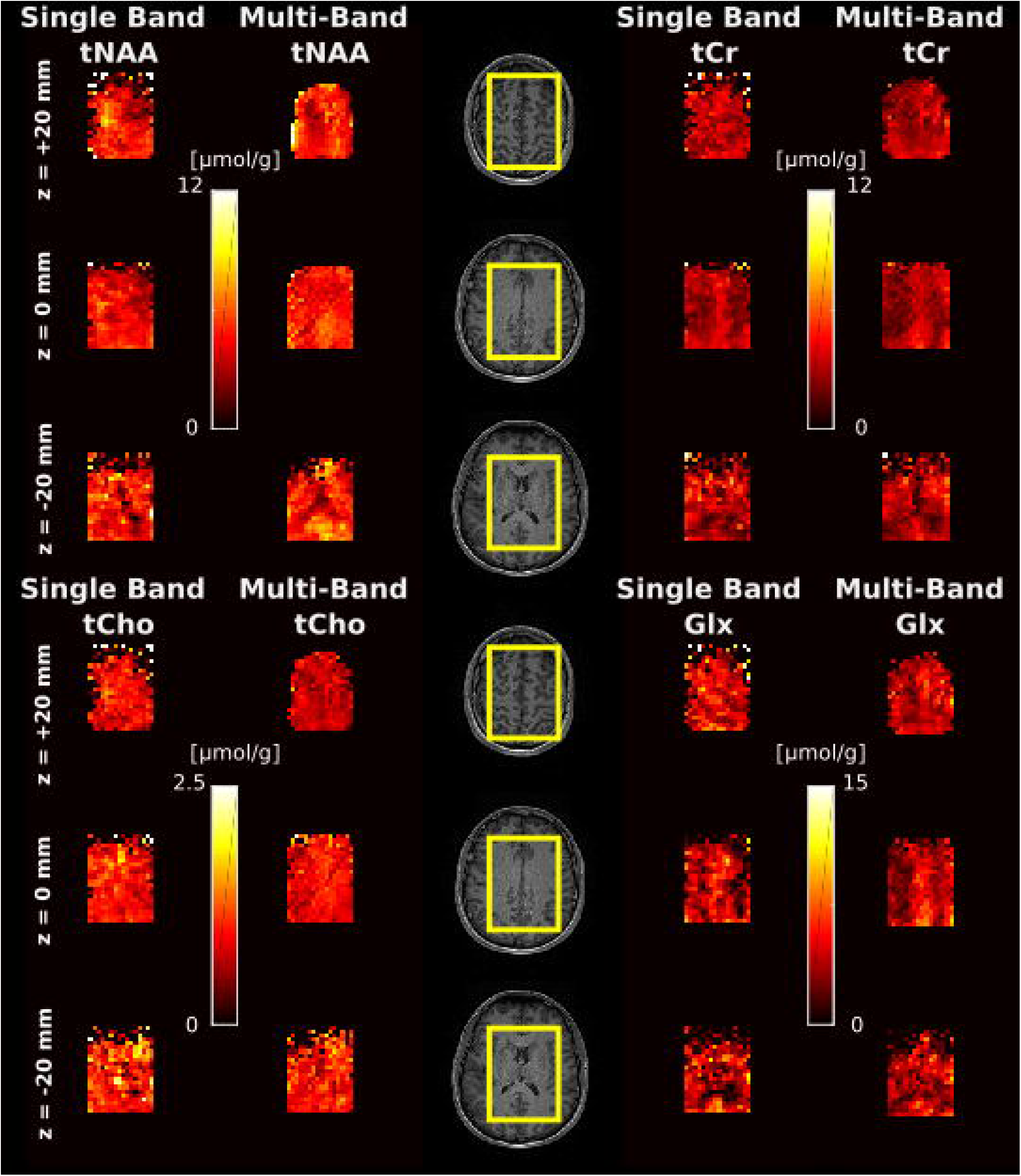
Metabolite distribution maps obtained with single and multi-band acquisitions from a subject. Absolute metabolite concentration maps from a nominal voxel dimension of 0.25 mL for tNAA, tCr, tCho, Glx illustrated with corresponding anatomical locations.

**Figure 8.**
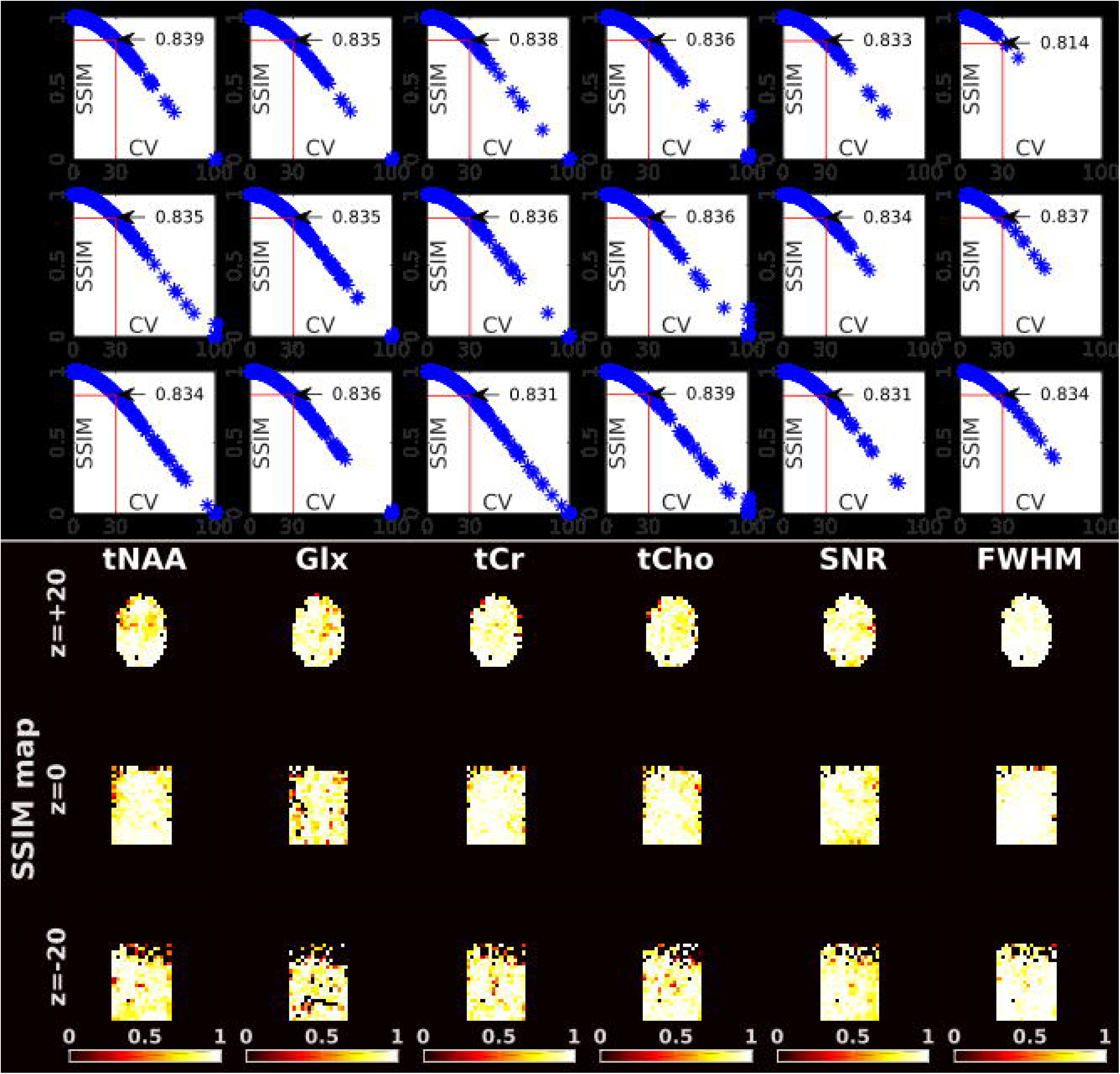
Structural similarity (SSIM) quality maps between multi-band and single-band acquisitions were were illustrated for tNAA, tCr, SNR and FWHM as estimated by LCModel. Comparison of the SSIM index and coefficient of variance (CV) between multi-and single-band acquisitions were also illustrated. A CV of 30 % and its corresponding SSIM index for each map were highlighted with a black arrow.

## Discussion

In this work, we demonstrated a non-water suppressed multi-band MRSI acquisition using DW-CRT. The results from structural and multi-compartment metabolite phantoms show that single and multi-band acquisition schemes produce similar quality spectra. In line with the phantom measurement, the in vivo results exhibited multi-band acquisition spectroscopic images of NAA, tCho, tCr, and Glx with an in-plane resolution of 5×5 mm^2^ and a slice thickness of 10 mm. The metabolite concentration values measured using the short-TE MRSI of all slices for different tissues were consistent with the literature (see below).

Compared to the EPI base acceleration techniques, DW-CRT has been demonstrated to offer a substantial improvement in SNR (12). The present study further accelerates the DW-CRT with simultaneous multi-slice excitation combined with SENSE parallel imaging reconstruction. It had been demonstrated that z-blips using concentric rings result in less overlap and, therefore, better g-factors (21). In line with this previous study, using SENSE resulted in an average g-factor of slightly more than 1 using z-blipped DW-concentric rings.

Structural and multi-compartment phantom experiment findings were further supported by in vivo experiments. In agreement with phantom experiments, multi and single-band DW-CRTs resulted in similar metabolite concentrations, SNRs FWHMs as estimated by LCModel and their distributions as determined by SSIM and CV analysis.

For the multi-band MRSI acquisition with a resolution of 0.25 mL, the achieved in vivo spectral quality allowed reliable quantification of primary brain metabolites with a CRLB of less than 50% using LCModel analysis. Concentration distributions of metabolites quantified in this study were in good agreement with previously reported values acquired from the same brain locations and revealed significant variations between brain tissues. tNAA and tCr have low values in the regions close to the skull, nasal cavity, and ventricles. Glx and tCr showed higher concentrations in the region of gray matter compared with white matter. In addition, we found elevated tCho, specifically tCho/tCr, in WM in comparison with GM.

There remain several limitations of the implemented methods. In this study, we chose to utilize STEAM localization with a penalty of two-fold signal loss compared with PRESS or semi-LASER since short TE values can easily be achieved with lower peak RF pulse powers, yielding a lower specific absorption rate (SAR). The use of the lipid penalty may also affect the quality of the absolute metabolite measures across the acquired region and reduce sensitivity to some metabolites. Using outer volume suppression pulses to suppress the lipid contamination during acquisition could make lipid removal unnecessary. This acquisition strategy should be explored in future studies.

In conclusion, here we show that MB DW-CRT acquisition in combination with non-water suppressed metabolite-cycled STEAM pulse localization allows fast, high-resolution 3D-MRSI to be acquired at 3 Tesla, with essentially no spatial or spectral penalty. Given the wide availability of 3 Tesla MRI scanners, this approach may have wide application in both clinical and research settings.

