## Supporting Table 1 for "Density-Weighted Concentric Ring Trajectory using simultaneous multi-band acceleration: 3D Metabolite-cycled Magnetic Resonance Spectroscopy Imaging at 3 T"

**Supporting Table 1 The number of voxels with a CV <30% between single- and multi-band acquisitions of phantom measurement for tNAA, tCr, SNR and FWHM as estimated by LCModel.**

| PHANTOM | Slice | tNAA | tCr | SNR | FWHM |
| --- | --- | --- | --- | --- | --- |
| Percentage of CV<30 | Z=20 mm | 434/677=64.1% | 768/911=84.3% | 798/916=87.1% | 656/725=90.5% |
|  | Z=0 mm | 409/744=55% | 773/915=84.5% | 733/915=80.1% | 621/693=89.6% |
|  | Z= -20 mm | 502/764=65.7% | 723/932=77.6% | 724/935=77.4% | 688/767=89.7% |

**Supporting Table 2 The number of voxels with a CV <30% between single- and multi-band acquisitions of all in-vivo subjects for tNAA, tCr, SNR and FWHM as estimated by LCModel.**

| IN-VIVO | Subject | Slice | tNAA | Glx | tCr | tCho | SNR | FWHM |
| --- | --- | --- | --- | --- | --- | --- | --- | --- |
| Percentage of CV<30 | NO1 | Z=20 mm | 265/293=90.4% | 193/277=69.7% | 262/294=89.1% | 260/294=88.4% | 273/293=93.2% | 242/250=96.8% |
|  |  | Z=0 mm | 305/333=91.6% | 206/290=71% | 287/334=85.9% | 298/327=91.1% | 299/333=89.8% | 239/247=96.8% |
|  |  | Z= -20 mm | 227/306=74.2% | 165/266=62% | 214/308=69.5% | 232/298=77.9% | 239/306=78.1% | 241/260=92.7% |
|  | NO2 | Z=20 mm | 141/171=82.5% | 96/163=58.9% | 139/164=84.8% | 125/163=76.7% | 138/171=80.7% | 119/130=91.5% |
|  |  | Z=0 mm | 276/334=82.6% | 212/317=66.9% | 282/321=87.9% | 270/319=84.6% | 223/334=66.8% | 238/252=94.4% |
|  |  | Z= -20 mm | 213/308=69.2% | 167/262=63.7% | 236/312=75.6% | 227/302=75.2% | 240/308=77.9% | 249/277=89.9% |
|  | NO3 | Z=20 mm | 232/272=85.3% | 202/261=77.4% | 246/270=91.1% | 252/268=94% | 194/272=71.3% | 223/231=96.5% |
|  |  | Z=0 mm | 275/337=81.6% | 162/267=60.7% | 280/334=83.8% | 276/324=85.2% | 190/337=56.4% | 271/286=94.8% |
|  |  | Z= -20 mm | 211/288=73.3% | 150/259=57.9% | 229/303=75.6% | 204/289=70.6% | 190/288=66% | 221/240=92.1% |
|  | NO4 | Z=20 mm | 250/292=85.6% | 207/274=75.5% | 257/292=88% | 247/277=89.2% | 269/292=92.1% | 230/240=95.8% |
|  |  | Z=0 mm | 271/331=81.9% | 183/318=57.5% | 270/335=80.6% | 268/326=82.2% | 252/331=76.1% | 253/271=93.4% |
|  |  | Z= -20 mm | 211/319=66.1% | 166/280=59.3% | 231/322=71.7% | 202/301=67.1% | 223/319=69.9% | 239/279=85.7% |
|  | NO5 | Z=20 mm | 163/219=74.4% | 170/221=76.9% | 191/222=86% | 179/222=80.6% | 186/219=84.9% | 175/178=98.3% |
|  |  | Z=0 mm | 273/328=83.2% | 211/309=68.3% | 282/327=86.2% | 269/326=82.5% | 280/328=85.4% | 266/276=96.4% |
|  |  | Z= -20 mm | 217/305=71.1% | 174/257=67.7% | 224/315=71.1% | 229/305=75.1% | 239/305=78.4% | 229/252=90.9% |
